## supplemental tables for "High-frequency synthetic apomixis in hybrid rice"

| T-DNA constructs | Co-cultivated calli | Number of HygR cell lines / co-cultivated callus | Resistant cell lines | Regeneration frequency | T0 plants |
|---|---|---|---|---|---|
| **T313 : sgMiMe** | 22 | 3.9 +/- 3.8 | 85 | 57.9 | 41 |
| **T314 : sg MiMe _pAtECS:BBM1** | 16 | 2.2 +/- 1 | 37 | 70.3 | |
| | 32 | 4.9 +/- 3.9 | 157 | 73.8 | 49 |
| **T315 : sgMiMe _pOsECS:BBM1** | 25 | 3.7 +/- 3.7 | 95 | 61.0 | |
| | 38 | 7.9 +/- 7.3 | 286 | 56.2 | 88 |

**Supplementary Table 1: Transformation efficiency in BRS-CIRAD 302 following co-cultivation of mature seed embryo-derived embryogenic calluses with *Agrobacterium* EHA105 suspensions carrying either the sgMiMe (T313), the sgMiMe_pAtECS:BBM1 (T314) or the sgMiMe_pOsECS:BBM1 (T315) binary plasmid.** Several hygromycin-resistant cell lines were formed per co-cultivated callus. The total number of hygromycin-resistant cell lines is shown: Only a sub-fraction of these lines was tested for regeneration. The number of T0 plants refers to the number of confirmed primary transformants transferred to the greenhouse.

| T-DNA construct | Biallelic | Homo-zygous | Monoallelic /WT | WT | Total plants | Edited | Editing efficiency | Number of fertile T0 plants | % Fertile plants |
|---|---|---|---|---|---|---|---|---|---|
| **T313 : sgMiMe** | 21 | 14 | 1 | 5 | 41 | 35 | 85,4 | 20 | 52.6 |
| **T314 : sg MiMe _pDD45:BBM1** | 27 | 13 | 0 | 9 | 49 | 40 | 81,6 | 33 | 49.3 |
| **T315 : sgMiMe _pECS:BBM1** | 42 | 22 | 5 | 19 | 88 | 64 | 72,7 | 36 | 43.9 |
| **Total** | | | | | 178 | 139 | 78,1 | 89 | 50 |

**Supplementary Table 2: *OsOSD1* editing efficiencies in primary (T0) transformants of BRS-CIRAD 302 harboring either the sgMiMe (T313), the sgMiMe_pAtOCS:BBM1 (T314) or the sgMiMe_pOsECS:BBM1 (T315) T-DNA.** The lesions observed can be homozygous (same alteration at the two alleles, biallelic (two distinct alterations at the two alleles) or heterozygous (or monoallelic: one allele altered, the other being wild-type (WT)). The number and frequency of fertile plants for each T-DNA construct is shown. The distribution of number of T1 seeds per fertile event is shown in Supplementary Figure 1.

| T0 Events | | Number of T-DNA copies | *Os OSD1* GTGAGAAATTCCGGCGGTAGGGCGGCGCTCGCCGACCCCTCGGGTGGTGGGTTCTTT | *PAIR1* CGCAGTCGCAGTTCTCGCAGGTCTCCCTCGACGACAACCTCCTCACCCTCCTCCCTTCCCCC | *Os REC8* CGTGGGAGTGTGAGTAGGTGGTGTGGCGATCGTGTACGAGAGGAAGGTGAAGGCTCTGTA | T1 seeds |
|---|---|---|---|---|---|---|
| T313: sgMiMe | **T313 4.3** | > 2 | Homozygous ( +T) | Biallelic (+T/+A) | Bilallelic (+1 / -6) | 127 |
| | **T313 9.1** | 1 | Biallelic (+T/+A) | Biallelic (-2/+1) | Homozygous (-C) | 164 |
| | **T313 12.2** | > 2 | Biallelic (+1 / -5) | Homozygous (+A) | Biallelic (-3 +1) | 227 |
| | **T313 21** | 2 | Biallelic (-47+46 / +1) | Biallelic (+T/+A) | Biallelic (+1 / -5+1) | 168 |
| | **T313 22.1** | 1 | WT | WT | WT | 150 |
| | **T313 22.2** | 1 | WT | WT | WT | 200 |
| T314: sgMiMe _pAtECS:BBM1 | **T314 3.1** | 1 | WT | WT | WT | 59 |
| | **T314 12.2** | 2 | Biallelic (-1/+1) | Biallelic (+T/+A) | Homozygous (+G) | 68 |
| | **T314 12.3** | 1 | WT | WT | WT | 664 |
| | **T314 15.1** | > 2 | Biallelic (+T/+AG) | Homozygous (+A) | Homozygous (-5+1 ) | 182 |
| | **T314 15.3** | > 2 | He (WT/+2) | Homozygous (+A) | Homozygous (--4) | 136 |
| | **T314 16** | 2 | Biallelic (+1 / -8+3) | Homozygous (+A) | Biallelic (-3+G / -2+G) | 91 |
| | **T314 23.1** | 1 | Biallelic (+2 / +1) | Biallelic (+G/+A) | Biallelic (+1 / -24) | 31 |
| | **T314 37.7** | 2 | Biallelic (+1/-5) | Homozygous (+T) | Biallelic (-3/+G) | 82 |
| | **T314 37.9** | 1 | Homozygous (-7) | Biallelic (-2/+1) | Biallelic (-6 / -17) | 14 |
| | **T314 44.1** | 2 | Biallelic (-7 / +1) | Homozygous (+A) | Biallelic (+1 / -2+1) | 15 |
| | **T314 44.2** | 2 | Biallelic (-7 / +1) | Homozygous (+A) | Biallelic (+1 / -6+1) | 33 |
| | **T314 46.2** | 1 | Biallelic (-7 / +1) | Homozygous (+A) | Biallelic (+1 / -5+1) | 52 |
| T315: sgMiMe _pOsECS:BBM1 | **T315 1.1** | > 2 | Homozygous (+2) | Biallelic(-2/+1) | Biallelic (+1 / -5+1) | 14 |
| | **T315 3.2** | 1 | Biallelic (-22 / -34+4) | Homozygous (-28) | Biallelic (-3/-4) | 141 |
| | **T315 3.3** | 1 | Biallelic (-4/-22) | Homozygous (+T) | Bilallelic (-26/+1) | 182 |
| | **T315 5.1** | > 2 | Homozygous (indel : TA/ AG -19) | Biallelic (+1 / -3) | Homozygous (-5+1) | 14 |
| | **T315 5.4** | 1 | Biallelic (+T/+A) | Biallelic (+G/+T) | Biallelic (+1 / -5) | 114 |
| | **T315 5.5** | 1 | Biallelic (+T/-22) | Biallelic (+G/+T) | Biallelic (+1 / -6+1) | 64 |
| | **T315 6.1** | 2 | Biallelic (+1 / -1) | Heterozygous (WT / -12) | Heterozygous (WT / -35) | 86 |
| | **T315 7.2** | 2 | Biallelic (-19/+14/ del) | Biallelic (+C/+A) | Biallelic (-24 / +1) | 150 |
| | **T315 7.4** | > 2 | Biallelic (-19+14 / -21+90) | Biallelic (+C/+A) | Biallelic (-4 / -24) | 73 |
| | **T315 8.1** | 1 | Homozygous (-5) | Homozygous (+G) | Homozygous (+C) | 264 |
| | **T315 8.2** | 1 | Biallelic (-5/+A) | Homozygous (+G) | Homozygous (+C) | 195 |
| | **T315 8.3** | 1 | Biallelic (-5/+A) | Homozygous (+G) | Homozygous (+C) | 123 |
| | **T315 14.1** | 1 | WT | WT | WT | 9 |
| | **T315 14.2** | 1 | WT | WT | WT | 23 |
| | **T315 14.3** | 1 | WT | WT | WT | 73 |
| | **T315 16.1** | > 2 | Biallelic (+1/-2) | Biallelic (-A/+A) | Biallelic(-15 / -5) | 18 |
| | **T315 16.3** | 2 | Homozygous (+1) | Biallelic (+T/+A) | Biallelic (-2 / -4) | 14 |
| | **T315 21.1** | 1 | WT | WT | WT | 55 |
| | **T315 24.1** | > 2 | Homozygous (-29) | Biallelic (-2/+1) | Biallelic (-4 / -4+1) | 50 |
| | **`T315 31.1** | > 2 | Biallelic (-39+2 / -7) | Homozygous (-A) | Biallelic (-6 / -6+3) | 24 |
| | **T315 31.4** | 1 | Biallelic (-2/+1) | Biallelic (+1/-7) | Biallelic (+1 / -24) | 36 |
| | **T315 41.2** | 1 | Biallelic (+T/+A) | Biallelic (+1 / -4/+3) | Biallelic (-5 / -5+1) | 30 |
| | **T315 41.6** | > 2 | Homozygous (+1) | Biallelic (+T/+A) | Biallelic (-4+1) | 25 |

**Supplementary Table 3: Lesions observed at CRISPR target regions (red) in *OsOSD1*, *PAIR1* and *OsREC8* coding sequences in fertile BRS-CIRAD 302 primary (T0) transformation events harboring the sgMiMe (T313) sgMiMe_pAtECS:BBM1 (T314) and sgMiMe_pOsECS:BBM1 (T315) T-DNAs.** Homozygous and biallelic lesions are highlighted by different green background colors. Alleles exhibiting inserted or deleted nucleotides that diverge from a multiple of three are indicative of frameshift mutations and are likely inactivated. Wild-type (WT) or heterozygous target regions appear on an orange background. The number of seeds harvested from T0 plants is shown. For T313, only the six T0 events exhibiting the highest number of T1 seeds were analyzed. For T314 and T315 only the T0 events showing more than nine T1 seeds were analyzed. The T314 15.3 line is heterozygous at the OsOSD1 locus and homozygous mutant at PAIR1 and OsREC8. Such a mutation configuration should lead to plant sterility. However, sequencing of T1 progeny plants proved that a late mutation (-31 nt deletion + 3nt insertion) eventually occurred in the primary transformant, altering the remaining intact allele of OsOSD1 after T0 leaf sample collection for DNA analysis. Line T315 6.1 is a putative knock-out in OsOSD1 only, which is known to be fertile (Mieulet et al., 2016).

| T0 event | T-DNA copy | T1 plant | 2n | 4n | % Diploids | Reminder % diploids in T1s |
|---|---|---|---|---|---|---|
| T314 15.1 | >2 | T314 15.1/4 | 45 | 2 | 95,7 | |
| | | T314 15.1/6 | 40 | 1 | 97,6 | |
| | | T314 15.1/8 | 40 | 0 | 100,0 | |
| | | T314 15.1/10 | 40 | 2 | 95,2 | |
| | | T314 15.1/11 | 45 | 2 | 95,7 | |
| | | | 210 | 7 | 96,8 | 98.5 |
| T314 37.7 | 2 | T314 37.7/11 | 42 | 6 | 87,5 | |
| | | T314 37.7/14 | 41 | 10 | 80,4 | |
| | | T314 37.7/15 | 40 | 2 | 95,2 | |
| | | T314 37.7/19 | 37 | 4 | 90,2 | |
| | | T314 37.7/20 | 44 | 2 | 95,7 | |
| | | | 204 | 24 | 89,5 | 92.1 |
| T315 3.2 | 1 | T315 3.2/485 | 38 | 6 | 86,4 | |
| | | T315 3.2/492 | 40 | 4 | 90,9 | |
| | | T315 3.2/501 | 40 | 4 | 90,9 | |
| | | T315 3.2/503 | 44 | 0 | 100,0 | |
| | | T315 3.2/504 | 43 | 1 | 97,7 | |
| | | | 205 | 15 | 93,2 | 95.7 |
| T315 5.4 | 1 | T315 5.4/411 | 43 | 1 | 97,7 | |
| | | T315 5.4/412 | 41 | 3 | 93,2 | |
| | | T315 5.4/417 | 44 | 2 | 95,7 | |
| | | T315 5.4/409 | 47 | 1 | 97,9 | |
| | | T315 5.4/414 | 60 | 6 | 90,9 | |
| | | | 235 | 13 | 94,8 | 93.2 |
| T315 8.1 | 1 | T315 8.1/329 | 33 | 7 | 82,5 | |
| | | T315 8.1/318 | 39 | 1 | 97,5 | |
| | | T315 8.1/292 | 40 | 0 | 100,0 | |
| | | T315 8.1/322 | 38 | 2 | 95,0 | |
| | | T315 8.1/313 | 51 | 5 | 91,1 | |
| | | | 201 | 15 | 93,1 | 95.4 |
| T315 8.2 | 1 | T315 8.2/207 | 50 | 10 | 83,3 | |
| | | T315 8.2/211 | 39 | 7 | 84,8 | |
| | | T315 8.2/212 | 54 | 10 | 84,4 | |
| | | T315 8.2/234 | 56 | 3 | 94,9 | |
| | | T315 8.2/350 | 40 | 4 | 90,9 | |
| | | | 239 | 34 | 87,5 | 97.2 |

**Supplementary Table 4: Detail of distribution of ploidy level among T2 progenies of five T1 plants of T314 (sgMiMe_pAtECS:BBM1) 15.1 and 37.7 events, and T315 (sgMiMe_pOsECS:BBM1) 3.2, 5.4, 8.1 and 8.2 events.** Frequency of diploid plants at the T1 generation is provided for reference.

| | 2n | 4n | % diploids |
|---|---|---|---|
| T314 15.1 / 11 / 7 | 98 | 2 | 98,0 |
| T314 15.1 / 6 / 10 | 99 | 1 | 99,0 |
| T314 15.1 / 8 / 7 | 99 | 1 | 99,0 |
| | | | 98,7 |
| T314 37.7 / 19 / 4 | 93 | 7 | 93,0 |
| T314 37.7 / 20 /2 | 92 | 8 | 92,0 |
| T314 37.7 / 11/ 6 | 96 | 4 | 96,0 |
| | | | 93,7 |
| T315 8.1/ 181 /4 | 94 | 6 | 94,0 |
| T315 8.1/ 182 /10 | 92 | 8 | 92,0 |
| T315 8.1 /186/3 | 92 | 8 | 92,0 |
| | | | 92,7 |
| T315 5.4/ 66/10 | 96 | 4 | 96,0 |
| T315 5.4/ 70/4 | 98 | 2 | 98,0 |
| T315 5.4/ 67/7 | 96 | 4 | 96,0 |
| | | | 96,7 |

**Supplementary Table 5: Detail of distribution of ploidy level among T3 progenies (n=100) of three randomly selected T2 plants of T314 (sgMiMe_pAtECS:BBM1) 15.1 and 37.7 events, and T315 (sgMiMe_pOsECS:BBM1) 5.4 and 8.1 and 8.1 events**

| T0 Event/T1 plant | T2/F1 plant number | Plant height | Number of tillers | n-1 Leaf length | n-1 Leaf width | Flag leaf length | Flag leaf width | Main panicle lentgh |
|---|---|---|---|---|---|---|---|---|
| T314 15.1/4 | 8 | 84.3 a | 12.9 a | 47.8 a | 1.4 a | 27.1 a | 1.6 a | 24.4 a |
| T314 15.1/6 | 8 | 83.9 a | 14.1 a | 49.9 a | 1.5 a | 28,1 a | 1.6 a | 24.3 a |
| T314 15.1/8 | 6 | 83.8 a | 15.5 a | 54.3 a | 1.1 a | 27.7 a | 1.4 a | 25.4 a |
| T314 15.1/10 | 5 | 81.0 a | 13.4 a | 42.9 a | 1.5 a | 31.6 a | 1,7 a | 24.1 a |
| T314 15.1/11 | 9 | 80.6 a | 16.0 a | 47.1 a | 1.3 a | 27.3 a | 1.5 a | 23.8 a |
| **T314 15.1 mean** | | **82.5** | **14.5** | **47.3** | **1.4** | **28.3** | **1.6** | **24.2** |
| T314 37.7/11 | 7 | 85.7 a | 15.4 a | 45.5 a | 1.4 a | 26.0 a | 1.5 a | 23.9 a |
| T314 37.7/14 | 8 | 85.4 a | 16.3 a | 45.4 a | 1.4 a | 27.6 a | 1.5 a | 23.3 a |
| T314 37.7/15 | 8 | 85.0 a | 14.1 a | 49.2 a | 1.4 a | 28.9 a | 1.5 a | 23.8 a |
| T314 37.7/19 | 8 | 82.9 a | 15.9 a | 45.9 a | 1.3 a | 26.2 a | 1.5 a | 23.6 a |
| T314 37.7/20 | 9 | 85.9 a | 14.2 a | 44.3 a | 1.4 a | 25.7 a | 1.4 a | 23.6 a |
| **T314 37.7 mean** | | **85.0** | **15.2** | **46.0** | **1.4** | **26.9** | **1.5** | **23.6** |
| T315 5.4/63 | 7 | 85.0 a | 15.4 a | 45.4 a | 1.5 a | 25.9 a | 1.5 a | 24.4 a |
| T315 5.4/66 | 5 | 87.4 a | 12.6 a | 45.6 a | 1.5 a | 26.4 a | 1.6 a | 24.5 a |
| T315 5.4/67 | 10 | 89.1 a | 13.8 a | 43.9 a | 1.5 a | 23.8 a | 1.6 a | 24.6 a |
| T315 5.4/70 | 8 | 87.3 a | 14.3 a | 48.1 a | 1.7 a | 27.1 a | 1.7 a | 24.6 a |
| T315 5.4/86 | 8 | 88.0 a | 14.6 a | 49.1 a | 1.6 a | 26.9 a | 1.8 a | 25.3 a |
| **T315 5.4 mean** | | **87.3** | **14.4** | **46.5** | **1.6** | **26.0** | **1.6** | **24.6** |
| T315 8.1/181 | 8 | 83.4 a | 14.0 a | 48.4 a | 1.4 a | 27.8 a | 1.6 a | 24.8 a |
| T315 8.1/182 | 8 | 82.1 a | 13.9 a | 47.0 a | 1.5 a | 27.5 a | 1.6 a | 24.4 a |
| T315 8.1/186 | 5 | 81.5 a | 14.2 a | 51.1 a | 1.6 a | 29.6 a | 1.7 a | 26.1 a |
| T315 8.1/188 | 5 | 86.8 a | 13.4 a | 48.0 a | 1.4 a | 28.0 a | 1.6 a | 26.2 a |
| T315 8.1/189 | 8 | 83.3 a | 13.5 a | 48.3 a | 1.5 a | 30.2 a | 1.6 a | 26.1 a |
| **T315 8.1 mean** | | **83.2** | **13.9** | **48.5** | **1.5** | **28.6** | **1.6** | **25.3** |
| **BRS-CIRAD 302 mean** | **15** | **84.8** | **13.4** | **50.1** | **1,3** | **31.9** | **1.6** | **24.5** |

**Supplementary Table 6: Detail of plant traits scored in T2 progenies of five T1 plants of T314 (sgMiMe_pAtECS:BBM1) 15.1 and 37.7 events and T315 (sgMiMe_pOsECS:BBM1) 5.4 and 8.1 events.** Variation across the five T2 progenies of each event was individually examined for statistical significance using a Kruskal-Wallis test. Numbers followed by the same letter are not statistically different at the $\alpha$-risk of 0.05.

| sgRNA | Target sequence **(PAM)** |
|---|---|
| sgRNA OsOSD1/1 | GAGAAATTCCGGCGGTAGGG **CGG** |
| sgRNA OsOSD1/2 | GCGCTCGCCGACCCCTCGGG **TGG** |
| sgRNA PAIR1 | GGTGAGGAGGTTGTCGTCGA **GGG** |
| sgRNA OsREC8 | GTGTGGCGATCGTGTACGAG **AGG** |

| **Primer** | **5'-3' Primer sequence** | **Comments** |
|---|---|---|
| OsOSD1-F<br>OsOSD1-R | TTACTTGGAAGAGGCAGGAGCC<br>ACCTTGACGACTGACGTGATGTC | Amplification of sgOsOSD1 target site |
| PAIR1-F<br>PAIR1-R | GTGGTGTGGTGTGTTCAGGAG<br>TGGAATCCCCAATCAGTAAGGCAC | Amplification of sgPAIR1 target site |
| OsREC8-F<br>OsREC8-R | GCACTAAGGCTCTCCGGAATTCTC<br>AATGGATCAAGGAGGAGGCACC | Amplification of sgOsREC8 target site |
| pEC1.2:BBM1-F<br>pEC1.2:BBM1-R | TTCCCTTTCCCACACGCTAC<br>AGAAGGAGGAGGAGGAGAAGC | Ascertaining presence of the pEC1.2:BBM1 cassette |
| pECA1:BBM1-F<br>pECA1:BBM1-R | ACCGTCAATCCTTTCCCATTCCTC<br>GAAGTCCTCCAGCTTCGGC | Ascertaining presence of the pECA1:BBM1 cassette |
| Cas9-F<br>Cas9-R | TGCCTGCGGAGGATAGCATGAAGCTC<br>TACCACGAGAAGTACCCGACCATCT | Ascertaining presence of Cas9 |
| HPT-F<br>HPT-R | CTGAACTCACCGCGACGTCTG<br>GGCGTCGGTTTCCACTATCG | Determination of T-DNA copy number integrated in T0 plants |
| RM1-F<br>RM1-R | GCGAAAACACAATGCAAAAA<br>GCGTTGGTTGGACCTGAC | Amplification of SSR locus RM1 located on chr 1 |
| RM25-F<br>RM25-R | GGAAAGAATGATCTTTTCATGG<br>CTACCATCAAAACCAATGTTC | Amplification of SSR locus RM25 located on chr 8 |
| RM215-F<br>RM215-R | CAAAATGGAGCAGCAAGAGC<br>TGAGCACCTCCTTCTCTGTAG | Amplification of SSR locus RM215 located on chr 9 |
| RM287-F<br>RM287-R | TTCCCTGTTAAGAGAGAAATC<br>GTGTATTTGGTGAAAGCAAC | Amplification of SSR locus RM287 located on chr 11 |

**Supplementary table 7: list of primers used for the molecular characterization of transformants and SSR genotyping.**
