## supplemental figures for "High-frequency synthetic apomixis in hybrid rice"

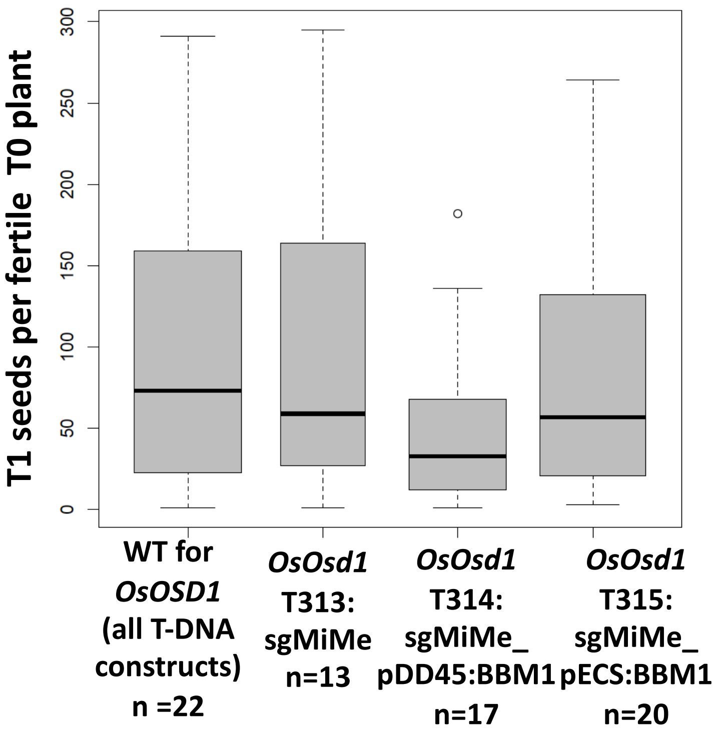


Supplementary figure 1

A

| | RM1 | | RM25 | | RM215 | | RM287 | |
|---|---|---|---|---|---|---|---|---|
| | Allele 1 | Allele 2 | Allele 1 | Allele 2 | Allele 1 | Allele 2 | Allele 1 | Allele 2 |
| 1F parent | 119 | 119 | 162 | 162 | 167 | 167 | 134 | 134 |
| 1F parent | 119 | 119 | 162 | 162 | 167 | 167 | 134 | 134 |
| D24 parent | 125 | 125 | 164 | 164 | 163 | 163 | 132 | 132 |
| D24 parent | 125 | 125 | 164 | 164 | 163 | 163 | 132 | 132 |
| BRS-CIRAD 302 | 119 | 125 | 162 | 164 | 163 | 167 | 132 | 134 |
| BRS-CIRAD 302 | 119 | 125 | 162 | 164 | 163 | 167 | 132 | 134 |
| BRS-CIRAD 302 | 119 | 125 | 162 | 164 | 163 | 167 | 132 | 134 |
| F2 progeny 1 | 119 | 125 | 162 | 164 | 163 | 167 | 132 | 134 |
| F2 progeny 2 | 119 | 125 | 162 | 164 | 163 | 167 | 132 | 134 |
| F2 progeny 3 | 119 | 125 | 162 | 164 | 163 | 163 | 132 | 134 |
| F2 progeny 4 | 119 | 119 | 162 | 164 | 163 | 167 | 134 | 134 |
| F2 progeny 5 | 119 | 125 | 162 | 164 | 163 | 167 | 134 | 134 |
| F2 progeny 6 | 119 | 119 | 162 | 164 | 163 | 167 | 134 | 134 |
| F2 progeny 7 | 125 | 125 | 164 | 164 | 163 | 163 | 134 | 134 |
| F2 progeny 8 | 119 | 125 | 164 | 164 | 167 | 167 | 134 | 134 |
| F2 progeny 9 | 119 | 125 | 164 | 164 | 163 | 167 | 134 | 134 |
| F2 progeny 10 | 119 | 125 | 162 | 162 | 167 | 167 | 132 | 132 |
| F2 progeny 11 | 119 | 119 | 162 | 162 | 167 | 167 | 134 | 134 |
| F2 progeny 12 | 119 | 125 | 164 | 164 | 163 | 163 | 132 | 132 |

B

| | | | | | | | | |
|---|---|---|---|---|---|---|---|---|
| T313 12.1 1 | 119 | 125 | 162 | 164 | 163 | 167 | 132 | 134 |
| T313 12.1 2 | 119 | 125 | 162 | 164 | 163 | 167 | 132 | 134 |
| T313 12.1 3 | 119 | 125 | 162 | 164 | 163 | 167 | 132 | 134 |
| T313 12.1 4 | 119 | 125 | 162 | 164 | 163 | 167 | 132 | 134 |
| T313 12.1 5 | 119 | 125 | 162 | 164 | 163 | 167 | 132 | 134 |
| T313 12.1 6 | 119 | 125 | 162 | 164 | 163 | 167 | 132 | 134 |
| T313 12.1 7 | 119 | 125 | 162 | 164 | 163 | 167 | 132 | 134 |
| T313 12.1 8 | 119 | 125 | 162 | 164 | 163 | 167 | 132 | 134 |
| T313 12.1 9 | 119 | 125 | 162 | 164 | 163 | 167 | 132 | 134 |
| T313 12.1 10 | 119 | 125 | 162 | 164 | 163 | 167 | 132 | 134 |
| T313 21 1 | 119 | 125 | 162 | 164 | 163 | 167 | 132 | 134 |
| T313 21 2 | 119 | 125 | 162 | 164 | 163 | 167 | 132 | 134 |
| T313 21 3 | 119 | 125 | 162 | 164 | 163 | 167 | 132 | 134 |
| T313 21 4 | 119 | 125 | 162 | 164 | 163 | 167 | 132 | 134 |
| T313 21 5 | 119 | 125 | 162 | 164 | 163 | 167 | 132 | 134 |
| T313 21 6 | 119 | 125 | 162 | 164 | 163 | 167 | 132 | 134 |
| T313 21 7 | 119 | 125 | 162 | 164 | 163 | 167 | 132 | 134 |
| T313 21 8 | 119 | 125 | 162 | 164 | 163 | 167 | 132 | 134 |
| T313 21 9 | 119 | 125 | 162 | 164 | 163 | 167 | 132 | 134 |
| T313 21 10 | 119 | 125 | 162 | 164 | 163 | 167 | 132 | 134 |

C

| | | | | | | | | |
|---|---|---|---|---|---|---|---|---|
| T314 15.1 1 | 119 | 125 | 162 | 164 | 163 | 167 | 132 | 134 |
| T314 15.1 2 | 119 | 125 | 162 | 164 | 163 | 167 | 132 | 134 |
| T314 15.1 3 | 119 | 125 | 162 | 164 | 163 | 167 | 132 | 134 |
| T314 15.1 4 | 119 | 125 | 162 | 164 | 163 | 167 | 132 | 134 |
| T314 15.1 5 | 119 | 125 | 162 | 164 | 163 | 167 | 132 | 134 |
| T314 15.1 6 | 119 | 125 | 162 | 164 | 163 | 167 | 132 | 134 |
| T314 15.1 7 | 119 | 125 | 162 | 164 | 163 | 167 | 132 | 134 |
| T314 15.1 8 | 119 | 125 | 162 | 164 | 163 | 167 | 132 | 134 |
| T314 15.1 9 | 119 | 125 | 162 | 164 | 163 | 167 | 132 | 134 |
| T314 15.1 10 | 119 | 125 | 162 | 164 | 163 | 167 | 132 | 134 |
| T314 15.1 11 | 119 | 125 | 162 | 164 | 163 | 167 | 132 | 134 |
| T314 15.1 12 | 119 | 125 | 162 | 164 | 163 | 167 | 132 | 134 |
| T314 15.1 13 | 119 | 125 | 162 | 164 | 163 | 167 | 132 | 134 |
| T314 15.1 14 | 119 | 125 | 162 | 164 | 163 | 167 | 132 | 134 |
| T314 15.1 15 | 119 | 125 | 162 | 164 | 163 | 167 | 132 | 134 |
| T314 15.1 16 | 119 | 125 | 162 | 164 | 163 | 167 | 132 | 134 |
| T314 15.1 17 | 119 | 125 | 162 | 164 | 163 | 167 | 132 | 134 |
| T314 15.1 18 | 119 | 125 | 162 | 164 | 163 | 167 | 132 | 134 |
| T314 15.1 19 | 119 | 125 | 162 | 164 | 163 | 167 | 132 | 134 |
| T314 15.1 20 | 119 | 125 | 162 | 164 | 163 | 167 | 132 | 134 |
| T314 15.1 21 | 119 | 125 | 162 | 164 | 163 | 167 | 132 | 134 |
| T314 15.1 22 | 119 | 125 | 162 | 164 | 163 | 167 | 132 | 134 |
| T314 15.1 23 | 119 | 125 | 162 | 164 | 163 | 167 | 132 | 134 |
| T314 15.1 24 | 119 | 125 | 162 | 164 | 163 | 167 | 132 | 134 |
| T314 15.1 25 | 119 | 125 | 162 | 164 | 163 | 167 | 132 | 134 |
| T314 15.1 26 | 119 | 125 | 162 | 164 | 163 | 167 | 132 | 134 |
| T314 15.1 27 | 119 | 125 | 162 | 164 | 163 | 167 | 132 | 134 |
| T314 15.1 28 | 119 | 125 | 162 | 164 | 163 | 167 | 132 | 134 |
| T314 15.1 29 | 119 | 125 | 162 | 164 | 163 | 167 | 132 | 134 |
| T314 15.1 30 | 119 | 125 | 162 | 164 | 163 | 167 | 132 | 134 |
| T314 15.1 31 | 119 | 125 | 162 | 164 | 163 | 167 | 132 | 134 |
| T314 15.1 32 | 119 | 125 | 162 | 164 | 163 | 167 | 132 | 134 |

D

| | | | | | | | | |
|---|---|---|---|---|---|---|---|---|
| T315_5.4_-63_B-_1 | 119 | 125 | 162 | 164 | 163 | 167 | 132 | 134 |
| T315_5.4_-63_B-_10 | 119 | 125 | 162 | 164 | 163 | 167 | 132 | 134 |
| T315_5.4_-63_B-_11 | 119 | 125 | 162 | 164 | 163 | 167 | 132 | 134 |
| T315_5.4_-63_B-_12 | 119 | 125 | 162 | 164 | 163 | 167 | 132 | 134 |
| T315_5.4_-63_B-_13 | 119 | 125 | 162 | 164 | 163 | 167 | 132 | 134 |
| T315_5.4_-63_B-_14 | 119 | 125 | 162 | 164 | 163 | 167 | 132 | 134 |
| T315_5.4_-63_B-_15 | 119 | 125 | 162 | 164 | 163 | 167 | 132 | 134 |
| T315_5.4_-63_B-_16 | 119 | 125 | 162 | 164 | 163 | 167 | 132 | 134 |
| T315_5.4_-63_B-_17 | 119 | 125 | 162 | 164 | 163 | 167 | 132 | 134 |
| T315_5.4_-63_B-_18 | 119 | 125 | 162 | 164 | 163 | 167 | 132 | 134 |
| T315_5.4_-63_B-_19 | 119 | 125 | 162 | 164 | 163 | 167 | 132 | 134 |
| T315_5.4_-63_B-_2 | 119 | 125 | 162 | 164 | 163 | 167 | 132 | 134 |
| T315_5.4_-63_B-_20 | 119 | 125 | 162 | 164 | 163 | 167 | 132 | 134 |
| T315_5.4_-63_B-_3 | 119 | 125 | 162 | 164 | 163 | 167 | 132 | 134 |
| T315_5.4_-63_B-_4 | 119 | 125 | 162 | 164 | 163 | 167 | 132 | 134 |
| T315_5.4_-63_B-_5 | 119 | 125 | 162 | 164 | 163 | 167 | 132 | 134 |
| T315_5.4_-63_B-_6 | 119 | 125 | 162 | 164 | 163 | 167 | 132 | 134 |
| T315_5.4_-63_B-_7 | 119 | 125 | 162 | 164 | 163 | 167 | 132 | 134 |
| T315_5.4_-63_B-_8 | 119 | 125 | 162 | 164 | 163 | 167 | 132 | 134 |
| T315_5.4_-63_B-_9 | 119 | 125 | 162 | 164 | 163 | 167 | 132 | 134 |
| T315_5.4_-66_B-_1 | 119 | 125 | 162 | 164 | 163 | 167 | 132 | 134 |
| T315_5.4_-66_B-_10 | 119 | 125 | 162 | 164 | 163 | 167 | 132 | 134 |
| T315_5.4_-66_B-_11 | 119 | 125 | 162 | 164 | 163 | 167 | 132 | 134 |
| T315_5.4_-66_B-_12 | 119 | 125 | 162 | 164 | 163 | 167 | 132 | 134 |
| T315_5.4_-66_B-_13 | 119 | 125 | 162 | 164 | 163 | 167 | 132 | 134 |
| T315_5.4_-66_B-_14 | 119 | 125 | 162 | 164 | 163 | 167 | 132 | 134 |
| T315_5.4_-66_B-_15 | 119 | 125 | 162 | 164 | 163 | 167 | 132 | 134 |
| T315_5.4_-66_B-_16 | 119 | 125 | 162 | 164 | 163 | 167 | 132 | 134 |
| T315_5.4_-66_B-_17 | 119 | 125 | 162 | 164 | 163 | 167 | 132 | 134 |
| T315_5.4_-66_B-_18 | 119 | 125 | 162 | 164 | 163 | 167 | 132 | 134 |
| T315_5.4_-66_B-_19 | 119 | 125 | 162 | 164 | 163 | 167 | 132 | 134 |
| T315_5.4_-66_B-_2 | 119 | 125 | 162 | 164 | 163 | 167 | 132 | 134 |
| T315_5.4_-66_B-_20 | 119 | 125 | 162 | 164 | 163 | 167 | 132 | 134 |
| T315_5.4_-66_B-_21 | 119 | 125 | 162 | 164 | 163 | 167 | 132 | 134 |
| T315_5.4_-66_B-_22 | 119 | 125 | 162 | 164 | 163 | 167 | 132 | 134 |
| T315_5.4_-66_B-_3 | 119 | 125 | 162 | 164 | 163 | 167 | 132 | 134 |
| T315_5.4_-66_B-_4 | 119 | 125 | 162 | 164 | 163 | 167 | 132 | 134 |
| T315_5.4_-66_B-_5 | 119 | 125 | 162 | 164 | 163 | 167 | 132 | 134 |
| T315_5.4_-66_B-_6 | 119 | 125 | 162 | 164 | 163 | 167 | 132 | 134 |
| T315_5.4_-66_B-_7 | 119 | 125 | 162 | 164 | 163 | 167 | 132 | 134 |
| T315_5.4_-66_B-_8 | 119 | 125 | 162 | 164 | 163 | 167 | 132 | 134 |
| T315_5.4_-66_B-_9 | 119 | 125 | 162 | 164 | 163 | 167 | 132 | 134 |
| T315_5.4_-67_B-_1 | 119 | 125 | 162 | 164 | 163 | 167 | 132 | 134 |
| T315_5.4_-67_B-_10 | 119 | 125 | 162 | 164 | 163 | 167 | 132 | 134 |
| T315_5.4_-67_B-_11 | 119 | 125 | 162 | 164 | 163 | 167 | 132 | 134 |
| T315_5.4_-67_B-_12 | 119 | 125 | 162 | 164 | 163 | 167 | 132 | 134 |
| T315_5.4_-67_B-_13 | 119 | 125 | 162 | 164 | 163 | 167 | 132 | 134 |
| T315_5.4_-67_B-_14 | 119 | 125 | 162 | 164 | 163 | 167 | 132 | 134 |
| T315_5.4_-67_B-_15 | 119 | 125 | 162 | 164 | 163 | 167 | 132 | 134 |
| T315_5.4_-67_B-_16 | 119 | 125 | 162 | 164 | 163 | 167 | 132 | 134 |
| T315_5.4_-67_B-_17 | 119 | 125 | 162 | 164 | 163 | 167 | 132 | 134 |
| T315_5.4_-67_B-_18 | 119 | 125 | 162 | 164 | 163 | 167 | 132 | 134 |
| T315_5.4_-67_B-_19 | 119 | 125 | 162 | 164 | 163 | 167 | 132 | 134 |
| T315_5.4_-67_B-_2 | 119 | 125 | 162 | 164 | 163 | 167 | 132 | 134 |
| T315_5.4_-67_B-_20 | 119 | 125 | 162 | 164 | 163 | 167 | 132 | 134 |
| T315_5.4_-67_B-_22 | 119 | 125 | 162 | 164 | 163 | 167 | 132 | 134 |
| T315_5.4_-67_B-_23 | 119 | 125 | 162 | 164 | 163 | 167 | 132 | 134 |
| T315_5.4_-67_B-_24 | 119 | 125 | 162 | 164 | 163 | 167 | 132 | 134 |
| T315_5.4_-67_B-_25 | 119 | 125 | 162 | 164 | 163 | 167 | 132 | 134 |
| T315_5.4_-67_B-_3 | 119 | 125 | 162 | 164 | 163 | 167 | 132 | 134 |
| T315_5.4_-67_B-_4 | 119 | 125 | 162 | 164 | 163 | 167 | 132 | 134 |
| T315_5.4_-67_B-_5 | 119 | 125 | 162 | 164 | 163 | 167 | 132 | 134 |
| T315_5.4_-67_B-_7 | 119 | 125 | 162 | 164 | 163 | 167 | 132 | 134 |
| T315_5.4_-67_B-_8 | 119 | 125 | 162 | 164 | 163 | 167 | 132 | 134 |
| T315_5.4_-67_B-_9 | 119 | 125 | 162 | 164 | 163 | 167 | 132 | 134 |
| T315_5.4_-70_B-_1 | 119 | 125 | 162 | 164 | 163 | 167 | 132 | 134 |
| T315_5.4_-70_B-_10 | 119 | 125 | 162 | 164 | 163 | 167 | 132 | 134 |
| T315_5.4_-70_B-_11 | 119 | 125 | 162 | 164 | 163 | 167 | 132 | 134 |
| T315_5.4_-70_B-_12 | 119 | 125 | 162 | 164 | 163 | 167 | 132 | 134 |
| T315_5.4_-70_B-_13 | 119 | 125 | 162 | 164 | 163 | 167 | 132 | 134 |
| T315_5.4_-70_B-_14 | 119 | 125 | 162 | 164 | 163 | 167 | 132 | 134 |
| T315_5.4_-70_B-_15 | 119 | 125 | 162 | 164 | 163 | 167 | 132 | 134 |
| T315_5.4_-70_B-_16 | 119 | 125 | 162 | 164 | 163 | 167 | 132 | 134 |
| T315_5.4_-70_B-_17 | 119 | 125 | 162 | 164 | 163 | 167 | 132 | 134 |
| T315_5.4_-70_B-_18 | 119 | 125 | 162 | 164 | 163 | 167 | 132 | 134 |
| T315_5.4_-70_B-_19 | 119 | 125 | 162 | 164 | 163 | 167 | 132 | 134 |
| T315_5.4_-70_B-_2 | 119 | 125 | 162 | 164 | 163 | 167 | 132 | 134 |
| T315_5.4_-70_B-_20 | 119 | 125 | 162 | 164 | 163 | 167 | 132 | 134 |
| T315_5.4_-70_B-_21 | 119 | 125 | 162 | 164 | 163 | 167 | 132 | 134 |
| T315_5.4_-70_B-_22 | 119 | 125 | 162 | 164 | 163 | 167 | 132 | 134 |
| T315_5.4_-70_B-_3 | 119 | 125 | 162 | 164 | 163 | 167 | 132 | 134 |
| T315_5.4_-70_B-_4 | 119 | 125 | 162 | 164 | 163 | 167 | 132 | 134 |
| T315_5.4_-70_B-_5 | 119 | 125 | 162 | 164 | 163 | 167 | 132 | 134 |
| T315_5.4_-70_B-_6 | 119 | 125 | 162 | 164 | 163 | 167 | 132 | 134 |
| T315_5.4_-70_B-_7 | 119 | 125 | 162 | 164 | 163 | 167 | 132 | 134 |
| T315_5.4_-70_B-_8 | 119 | 125 | 162 | 164 | 163 | 167 | 132 | 134 |
| T315_5.4_-70_B-_9 | 119 | 125 | 162 | 164 | 163 | 167 | 132 | 134 |
| T315_5.4_-86_B-_1 | 119 | 125 | 162 | 164 | 163 | 167 | 132 | 134 |
| T315_5.4_-86_B-_10 | 119 | 125 | 162 | 164 | 163 | 167 | 132 | 134 |
| T315_5.4_-86_B-_11 | 119 | 125 | 162 | 164 | 163 | 167 | 132 | 134 |
| T315_5.4_-86_B-_12 | 119 | 125 | 162 | 164 | 163 | 167 | 132 | 134 |
| T315_5.4_-86_B-_13 | 119 | 125 | 162 | 164 | 163 | 167 | 132 | 134 |
| T315_5.4_-86_B-_14 | 119 | 125 | 162 | 164 | 163 | 167 | 132 | 134 |
| T315_5.4_-86_B-_15 | 119 | 125 | 162 | 164 | 163 | 167 | 132 | 134 |
| T315_5.4_-86_B-_16 | 119 | 125 | 162 | 164 | 163 | 167 | 132 | 134 |
| T315_5.4_-86_B-_17 | 119 | 125 | 162 | 164 | 163 | 167 | 132 | 134 |
| T315_5.4_-86_B-_18 | 119 | 125 | 162 | 164 | 163 | 167 | 132 | 134 |
| T315_5.4_-86_B-_19 | 119 | 125 | 162 | 164 | 163 | 167 | 132 | 134 |
| T315_5.4_-86_B-_2 | 119 | 125 | 162 | 164 | 163 | 167 | 132 | 134 |
| T315_5.4_-86_B-_20 | 119 | 125 | 162 | 164 | 163 | 167 | 132 | 134 |
| T315_5.4_-86_B-_21 | 119 | 125 | 162 | 164 | 163 | 167 | 132 | 134 |
| T315_5.4_-86_B-_22 | 119 | 125 | 162 | 164 | 163 | 167 | 132 | 134 |
| T315_5.4_-86_B-_23 | 119 | 125 | 162 | 164 | 163 | 167 | 132 | 134 |
| T315_5.4_-86_B-_24 | 119 | 125 | 162 | 164 | 163 | 167 | 132 | 134 |
| T315_5.4_-86_B-_25 | 119 | 125 | 162 | 164 | 163 | 167 | 132 | 134 |
| T315_5.4_-86_B-_26 | 119 | 125 | 162 | 164 | 163 | 167 | 132 | 134 |
| T315_5.4_-86_B-_3 | 119 | 125 | 162 | 164 | 163 | 167 | 132 | 134 |
| T315_5.4_-86_B-_4 | 119 | 125 | 162 | 164 | 163 | 167 | 132 | 134 |
| T315_5.4_-86_B-_5 | 119 | 125 | 162 | 164 | 163 | 167 | 132 | 134 |
| T315_5.4_-86_B-_6 | 119 | 125 | 162 | 164 | 163 | 167 | 132 | 134 |
| T315_5.4_-86_B-_7 | 119 | 125 | 162 | 164 | 163 | 167 | 132 | 134 |
| T315_5.4_-86_B-_8 | 119 | 125 | 162 | 164 | 163 | 167 | 132 | 134 |
| T315_5.4_-86_B-_9 | 119 | 125 | 162 | 164 | 163 | 167 | 132 | 134 |

Supplementary Figure 2

A

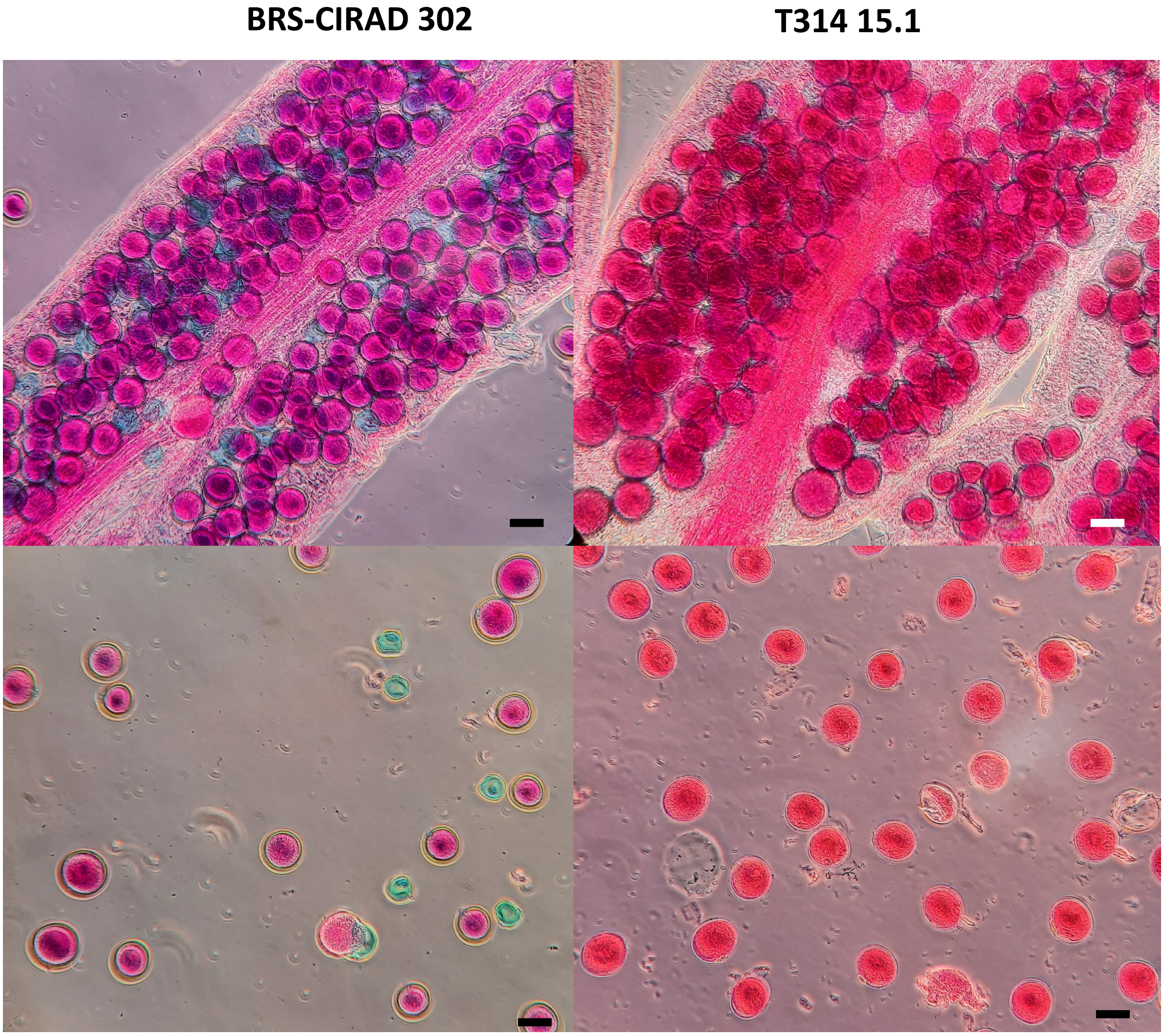


B

| Material | Plants | Average pollen viability | SD |
|---|---|---|---|
| BRS-CIRAD 302 | 4 | 59.1 | 7.0 |
| T2 T314 15.1 | 4 | 85.9 | 2.3 |
| T2 T314 37.7 | 6 | 95.3 | 1.7 |
| T2 T315 5.4 | 5 | 95.2 | 1.7 |
| T2 T315 8.1 | 4 | 94.6 | 1.9 |

Supplementary Figure 3

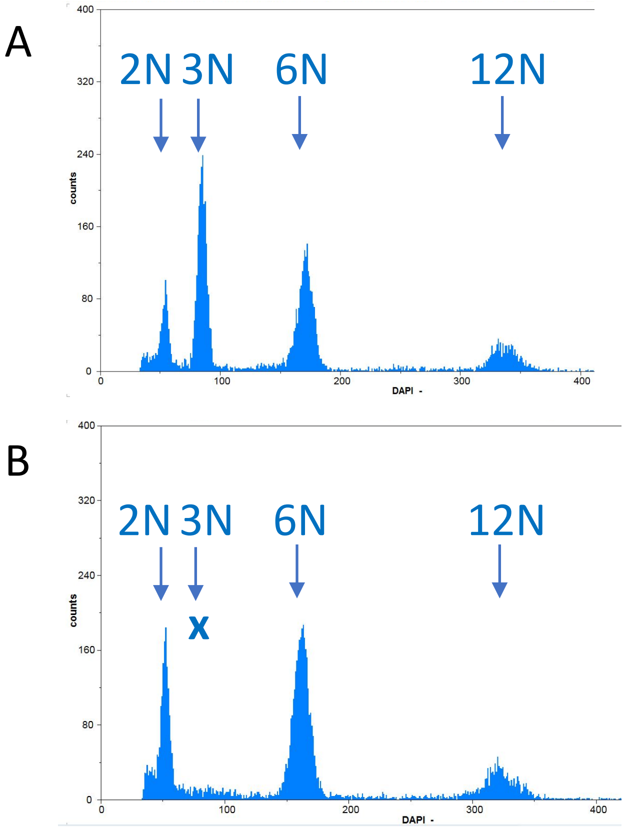


Supplementary Figure 4

A

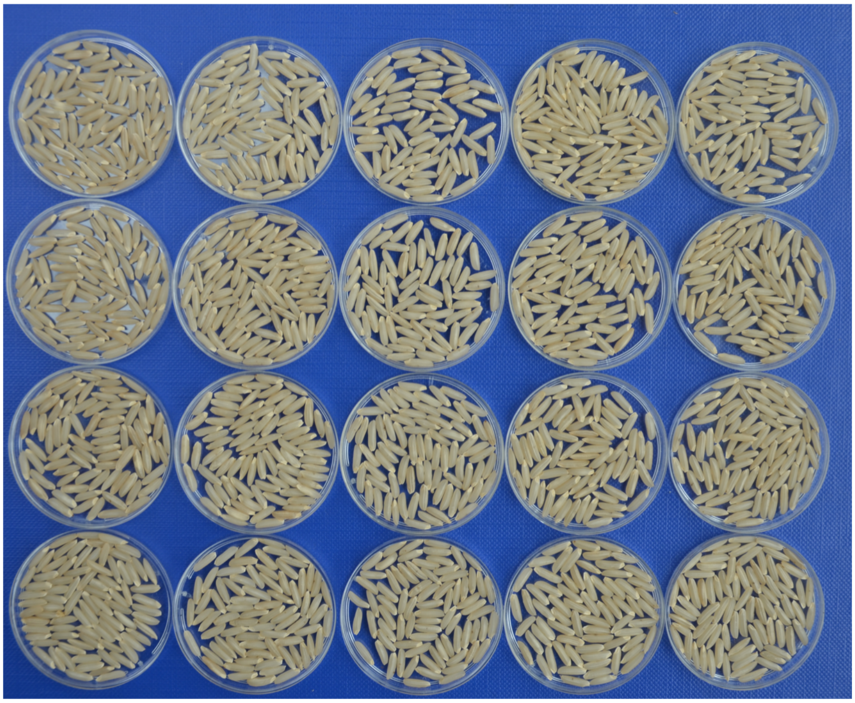


B

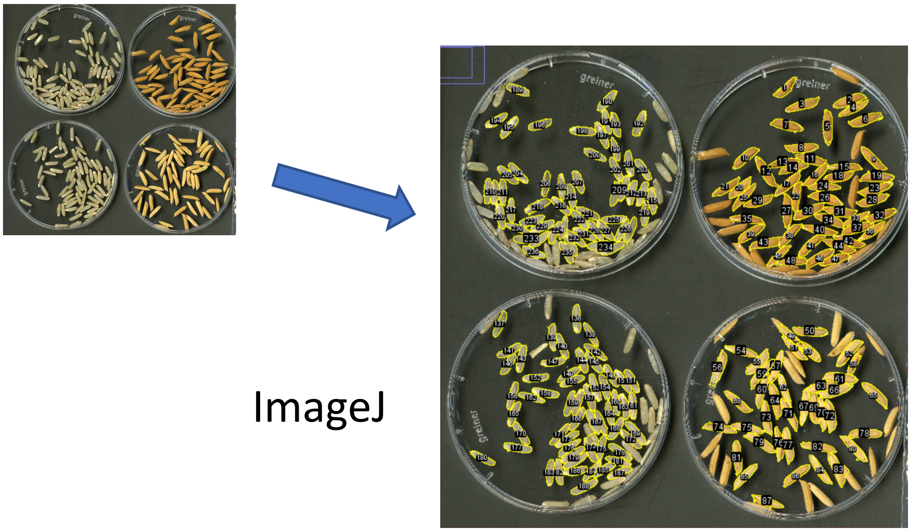


C

| | length (mm) | width (mm) | volume (mm3) | shape |
|---|---|---|---|---|
| 1F | 6.88 | 2.11 | 10.18 | 3.26 |
| D24 | 7.32 | 2.01 | 9.79 | 3.66 |
| BRS-CIRAD 302 | 7.17 | 2.19 | 11.36 | 3.28 |
| F2 seeds | 7.18 a,b | 2.11 a | 10.58 a,b | 3.42 a |
| T314 15.1 | 7.15 a,b | 2.05 a,b | 9.97 a,b | 3.50 a |
| T314 37.7 | 6.95 b | 2.02 b | 9.39 b | 3.45 a |
| T315 5.4 | 7.37 a | 2.12 a | 10.97 a | 3.48 a |
| T315 8.1 | 7.33 a | 2.08 a,b | 10.46 a,b | 3.54 a |
| Average apomictics | 7.18 | 2.06 | 10.13 | 3.49 |

Supplementary Figure 5
